# A benchmark of computational methods for correcting biases of established and unknown origin in CRISPR-Cas9 screening data

**DOI:** 10.1101/2024.01.30.577980

**Authors:** Alessandro Vinceti, Raffaele Iannuzzi, Isabella Boyle, Lucia Trastulla, Catarina D. Campbell, Francisca Vazquez, Joshua Dempster, Francesco Iorio

## Abstract

CRISPR-Cas9 dropout screens stand as formidable tools for investigating biology with unprecedented precision and scale. One of their principal applications involves probing large panels of immortalised human cancer cell lines for viability reduction responses upon systematic genetic knock-out at a genome-wide level, to identify novel cancer dependencies and therapeutic targets. However, biases in CRISPR-Cas9 screens’ data pose challenges, leading to potential confounding effects on their interpretation and compromising their overall quality. The mode of action of the Cas9 enzyme, exerted by the induction of DNA double-strand breaks at a locus targeted by a specifically designed single-guide RNA (sgRNA), is influenced by structural features of the target site, including copy number amplifications (CN bias). More worryingly, proximal targeted loci tend to generate similar gene-independent responses to CRISPR-Cas9 targeting (*proximity bias*), possibly due to Cas9-induced whole chromosome-arm truncations or other unknown genomic structural features and different chromatin accessibility levels.

Different computational methods have been proposed to correct these biases *in silico*, each based on different modelling assumptions. We have benchmarked seven of the latest methods, rigorously evaluating for the first time their ability to reduce both CN and proximity bias in the two largest publicly available cell-line-based CRISPR-Cas9 screens to date. We have also evaluated the capability of each method to preserve data quality and heterogeneity by assessing the extent to which the processed data allows accurate detection of true positive essential genes, established oncogenetic addictions, and known/novel biomarkers of cancer dependency.

Our analysis sheds light on the ability of each method to correct biases arising from structural properties and other possible unknown factors associated with CRISPR-Cas9 screen data under different scenarios. In particular, it shows that AC-Chronos outperforms other methods in correcting both CN and proximity biases when jointly processing multiple screens of models with available CN information, whereas CRISPRcleanR is the top performing method for individual screens or when CN information is not available for the screened models. In addition, Chronos and AC-Chronos yield a final dataset better able to recapitulate known sets of essential and non-essential genes.

Overall, our investigation provides guidance for the selection of the most appropriate bias-correction method, based on its strengths, weaknesses and experimental settings.

## Background

CRISPR-Cas9-mediated loss of function screens are revolutionising molecular biology and genomic research, offering an unparalleled ability to manipulate the DNA of model systems and living organisms with exquisite precision, ease and scale. This groundbreaking technology has empowered researchers to systematically explore the functions of individual genes on a genome-wide scale. Furthermore, it has opened up exciting possibilities for reconstructing gene-regulatory networks, delving deep into the mechanisms of human diseases, and uncovering novel therapeutic targets alongside their associated biomarkers, paving powerful new avenues towards the realisation of personalised medicine [1–7].

The CRISPR-Cas9 system is based on two components: the Cas9 enzyme and a single-guide RNA (sgRNA). The sgRNA binds to a specific region in the genome, where Cas9 is subsequently recruited, and guides the enzyme to induce DNA double-strand breaks (DSBs) at a specific genomic locus [8–11]. These DSBs trigger the cell to repair the DNA through non-homologous end joining, an error-prone mechanism that introduces small insertions/deletions. When this occurs within an intragenic locus, it gives rise to premature stop codons, effectively silencing the targeted gene [12].

One of the most promising applications of CRISPR-Cas9 technology has been screening large panels of immortalised human cancer cell lines. This is achieved through genome-wide libraries of single-guide RNAs (sgRNAs) used in a pooled fashion to systematically knock out individual genes, allowing for a thorough assessment of the resulting perturbations on cellular viability [3, 5, 6, 13]. This is quantified by the relative abundance of each individual sgRNA, which scars the genome of the receiving cells and can be used as a barcode for counting them via NGS sequencing of the final population of cells (expanded after a certain amount of time post-library transduction) [14].

Integrative analyses of the genetic dependency profiles derived from these screens have allowed identifying cancer vulnerabilities specific to different genomic contexts, contributing to the discovery of new anti-cancer therapeutic targets [7, 15, 16] and representing the core data source underlying a comprehensive cancer dependency map [17–22].

While the CRISPR-Cas9 technology holds significant translational potential, the robust detection of genes and pathways crucial for the viability of cancer cells through this tool requires particular attention [23].

Complications arise from the mechanism of action of the Cas9 enzyme itself, which is influenced by structural features of the targeted loci resulting in biases that can negatively impact the overall quality of the screens, hampering their interpretation and leading to false positives while identifying genes whose function is essential for cellular viability (essential or fitness genes).

In addition to sgRNA off-target effects resulting in unintended DNA cleavages and other hindering factors (e.g. heterogeneous guide on-target efficiency [24–26] and sub-optimal experimental conditions [27–29]), two primary factors related to the structural properties of the targeted genome have been associated with gene-independent responses to CRISPR-Cas9 targeting, hence identified as sources of bias in related data.

The first source of bias is genomic copy number (CN) amplifications [30–34], i.e. arising when a region targeted by a given sgRNA is present in multiple copies in the genome (**Fig. 1AB**). It has been demonstrated that this causes Cas9 inducing several DSBs which results in cell cycle arrest and cell death. This causes genes within CN amplified regions to be mistakenly identified as essential for cell viability (fitness genes) regardless of their function, expression and actual presence, as this effect is observed also when targeting CN amplified inter-genic loci [30–32]. Subsequently, we reported that biases observed in CRISPR-Cas9 screens arising from CN alterations are non-linear and heterogeneous, with variations between segments of differing CNs, and variations within segments of the same CNs (**Fig. 1CD**) [34]. This motivated a further study, which demonstrated that CN biases in CRISPR-Cas9 screens are specific to tandem and interspersed CN amplifications with intensities proportional to CN of the targeted gene/region relative to that of its hosting chromosome [33].

**Figure 1.**
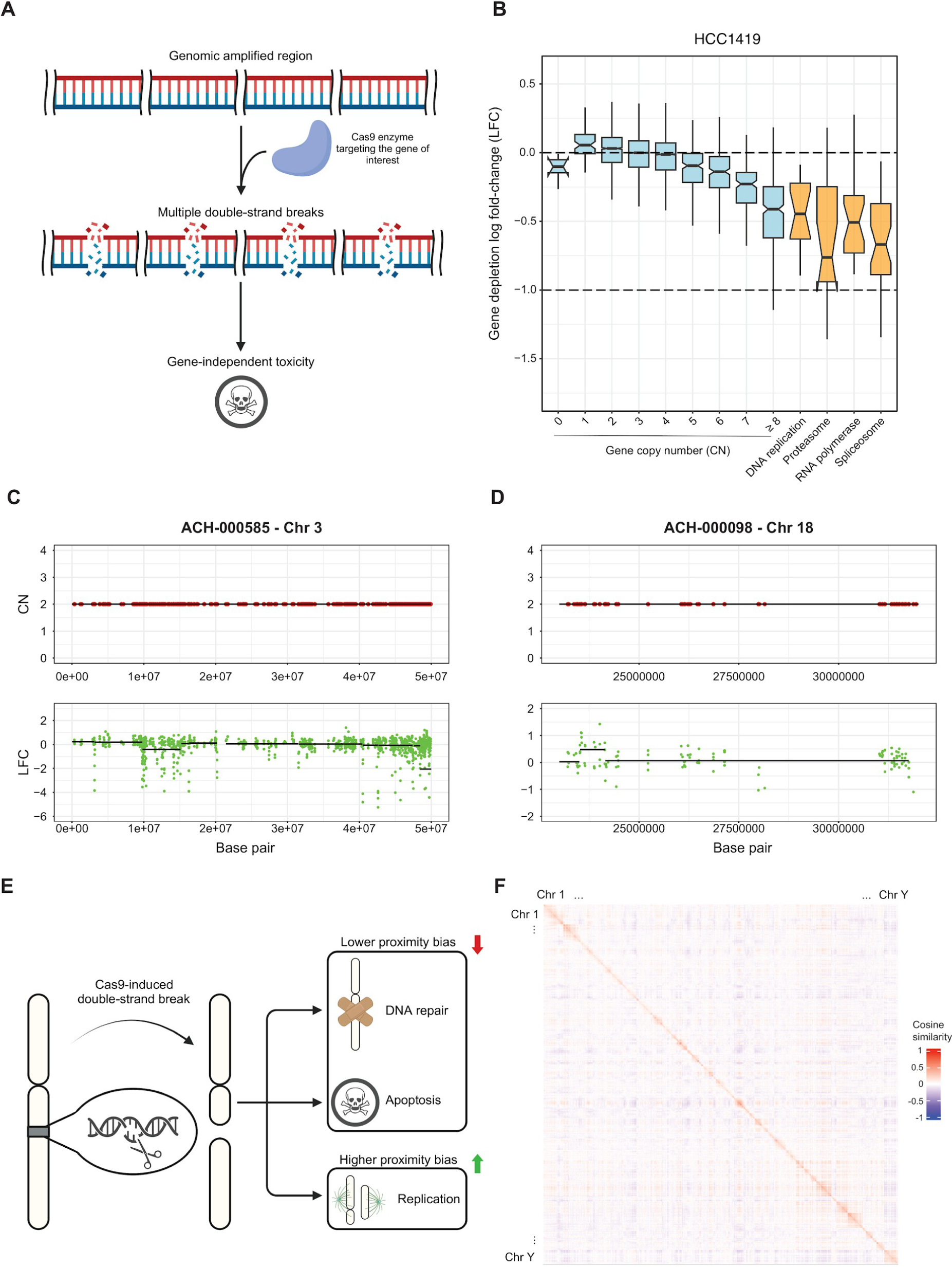
Overview of main structural features causing biases in CRISPR-Cas9 screens. A. Copy number amplification bias in CRISPR screening data: when the Cas9 enzyme targets a copy number amplified gene it induces multiple double-strand breaks (DSBs) resulting in a high cytotoxic effect independent of the gene’s function or expression. B. Gene-level depletion log fold change (LFC) values grouped by gene absolute copy number (in cyan) and biological functions (positive controls in orange) in the HCC1419 cell line. The positive controls are genes involved in housekeeping processes, thus characterised by strong essentiality for the viability of any cell (included for comparison). CD. Biased segments of gene depletion LFCs can be observed also within genomic segments of normal copy number and the bias can be even positive. C. A portion of chromosome 3 in the ACH-000585 cell line characterised by negatively biased segments of LFC within non-copy-number (CN) altered regions. D. A portion of chromosome 18 in the ACH-000098 cell line characterised by positively biased segments of depletion LFC within non-CN altered regions. E. Proximity bias in CRISPR screening data: when a functional p53 or other DNA-repairing factors are present, DSBs cause cell cycle arrest (until the genetic damage is fixed) or apoptosis. However, in the absence or reduced levels of functional p53, a DSB may not be repaired before mitosis. In this case, DNA segments originating from the break site may fail to segregate properly and get lost in future cell generations. The magnitude of this effect is strongly affected by the location of the gene along the chromosome arm. F. Pairwise cosine similarity between genes’ LFCs in the Project Achilles (v23Q2) dataset sorted by chromosome number across the human genome and based on genomic coordinates within the same chromosome, and quantile normalised to a normal distribution with mean 0 and standard deviation 0.2.

A second, more general source of bias, related to sets of proximal genes yielding suspiciously similar fitness effects when targeted with the CRISPR-Cas9 system (*proximity bias*), in an experiment-specific fashion and independently from their function/expression, was first reported by us [34] and exploited for the development of an unsupervised correction algorithm (CRISPRcleanR), which does not make any assumption on the bias cause. This proximity bias has been recently confirmed and linked to whole chromosome harm truncations resulting from the accumulation of DSBs in adjacent regions [35]. Furthermore, this has recently sparked concern in the community as Lazar and colleagues [35] have reported that the proximity bias is present and clearly visible in recent and widely used *cancer dependency map* datasets (such as the Broad Institute DepMap CRISPR 19Q3 and 22Q2), in which CRISPR-Cas9 induced viability phenotypes of genes located on the same chromosome arm are highly correlated, even when this data is post-processed with state-of-the-art tools (**Fig. 1EF**).

Lazar and colleagues [35] illustrate different scenarios that can be triggered by Cas9-induced DSBs within the cell: i) DNA repair with resulting restoration, ii) extensive damage leading to cell apoptosis, or iii) impairment of a biological function and cell mitosis with unrepaired DNA (**Fig. 1E**). The first two cases causes mild to no proximity bias, while the latter allows the cell to carry on its cycle at suboptimal rate upon the extension of the damage, resulting in a significantly stronger proximity bias. Many molecular investigations have suggested that large-scale chromosomal truncations occurring in a subset of cells might be responsible for this effect [36–38]. These truncations are responsible for the deletion of entire chromosomal regions that contain multiple genes, resulting in a mixed viability reduction profile given by the sum of the individual gene death phenotypes.

Several computational methods have been developed to correct biases in CRISPR-Cas9 screening data *in silico*, each based on different modelling assumptions.

Here, we present results from a comprehensive benchmark of seven of these methods, which are state-of-the-art and among the most widely used. We evaluate (for the first time, to the best of our knowledge) the ability of these methods to correct both CN and proximity biases in CRISPR-Cas9 screening data while minimising data distortion, ensuring that the potential of this type of data to unveil new therapeutic targets, human core-fitness genes, and cancer dependency biomarkers is not compromised. Finally, we provide guidance on selecting the most appropriate correction method, based on the investigative goal under consideration and the employed experimental settings.

## Results

### Overview of the benchmarked methods

We compared the performances of CRISPRcleanR (CCR) [34], Chronos [39], Crispy [33], Local Drop Out (LDO), Generalised Additive Model (GAM) [40], Model-based Analysis of Genome-wide CRISPR-Cas9 Knockout (MAGeCK) based on a maximum likelihood estimation method [41, 42], and the recent Geometric method introduced in [35], in terms of correction of biases observed in widely used large-scale CRISPR-Cas9 screening dataset. In addition, we included in our benchmark also Arm-correct Chronos (AC-Chronos): a new computational pipeline recently implemented to process the Broad Cancer Dependency Map [21] and used since its 23Q2 release (**Table 1**).

**Table 1.** List of benchmarked methods.

| Method | Input data format | Algorithm | Additional data required | Multiple or single-screen | Source |
| --- | --- | --- | --- | --- | --- |
| **Model-based Analysis of Genome-wide CRISPR-Cas9 Knockout (MAGeCK)** | FASTQ / BAM / sgRNA-level read counts | Supervised | Gene-level CN | Multi-screen | [41, 42] |
| **Chronos** | sgRNA-level read counts | Supervised | Gene-level CN | Multi-screen | [39] |
| **AC-Chronos** | sgRNA-level read counts | Supervised | Gene-level CN | Multi-screen | [39] |
| **Crispy** | sgRNA-level read counts | Supervised | Segment-level CN | Single-screen | [33] |
| **Generalised Additive Model (GAM)** | sgRNA-level read counts | Supervised | Gene-level CN | Single-screen | [40] |
| **Local Drop Out (LDO)** | sgRNA-level read counts | Unsupervised | None | Single-screen | [40] |
| **CRISPRcleanR (CCR)** | FASTQ / BAM / sgRNA-level read counts | Unsupervised | None | Single-screen | [34] |
| **Geometric** | sgRNA/gene-level LFC | Supervised | Basal Expression, Genomic coordinates | Single-screen | [35] |

Some of the methods included in our benchmark (CCR and LDO) work in an unsupervised way on the CRISPR-Cas9 screening data only, not requiring additional information about the screened model. Some other methods (Chronos, AC-Chronos, Crispy, GAM, MAGeCK, Geometric) work in a supervised way, requiring additional data derived from the basal characterisation of the screened model, typically its profile of CN alteration or its basal transcriptional profile.

Furthermore, some methods (CCR, Crispy, GAM, Geometric, LDO) are designed to process data from individual screens, whereas others (Chronos, AC-Chronos, MAGeCK) work on multiple screens in single executions, working on gene depletion signals assembled across multiple screened models to correct genes’ depletion scores in the individual screens.

The compared methods implement different algorithms, which are based on heterogeneous modelling assumptions that, in turn, build upon different hypotheses on the source of biases. For example, MAGeCK (Model-based Analysis of Genome-wide CRISPR/Cas9 Knockout) is a method that was originally designed to identify significantly positively and negatively selected sgRNAs (possibly collapsed on targeted gene-basis level) and associated statistical scores in genome-wide CRISPR-Cas9 screens [41, 42]. It employs a negative binomial distribution to model the overdispersion of CRISPR screening read counts, and then it applies a generalised linear model that accounts for sgRNA knockout efficiency and other complex experimental settings while determining positive/negative hits. In our analyses, we focused on its latest extension (MAGeCK MLE), which implements a maximum likelihood estimation framework [42]. This version also implements a CN bias correction by integrating gene-level CN data of the screened model as a covariate, optimising model parameters through an expectation-maximisation approach. The final beta scores represent gene fitness effects after bias correction, which can be interpreted as corrected gene depletion log fold changes (LFCs).

Chronos is a supervised method modelling the dynamics of cell population in CRISPR-Cas9 knockout experiments taking into account several parameters such as sgRNA efficiency, heterogeneous outcomes and the delay between the knockout and the emergence of the phenotype [39]. The method maximises the likelihood of the observed matrix of normalised read counts under a negative binomial distribution, like MAGeCK, from which gene effects are inferred. To remove CN bias, it constructs a two-dimensional cubic spline from the matrix of inferred gene effects and gene-level CN data. Like MAGeCK, Chronos also is designed to work on multiple screens. In AC-Chronos, gene fitness scores computed by Chronos across multiple CRISPR screens are further normalised within chromosome harms in each screen such that intra-arm median fitness effects in a screen matches the intra-arm median fitness effect across all processed screens (**Methods**).

Crispy utilises a squared-exponential kernel to model the impact of CN bias in CRISPR-Cas9 screens on a single-screen basis [33]. For each screen, the inputs are sgRNA read counts together with segment-level CN data. A Gaussian process regression is used to model nonlinearities between segment CN data and LFCs. After training, corrected LFCs are obtained by subtracting the estimated bias from the original LFCs. Similarly, GAM [40] is a supervised method that requires the CN profile of the screened model data as additional input. It incorporates nonlinear dependencies and it accounts for multiple predictor variables for bias correction. Both, Crispy- and GAM-corrected scores can be used in place of the original LFC scores and interpreted as corrected LFCs.

LDO [40] and CRISPRcleanR [34] are unsupervised methods for identifying genomic segments in which long lists of neighbouring genes display similar and strong drop-out values, suggestive of a bias which might or might be not due to CN amplification, i.e. these methods do not make any assumption on the source of observed bias.

LDO implements a regression tree to estimate this effect, which is then removed from the original LFC scores. CRISPRcleanR (CCR) applies a circular binary segmentation algorithm [43, 44] directly to genome-sorted sgRNA depletion LFCs across individual chromosomes in a cell line. The LFCs of the sgRNAs targeting genes in the detected seemingly biased genomic segments are then mean-centred, thus smoothing fitness effects that are putatively gene-independent.

The Geometric method, recently introduced by Lazar and colleagues [35], has been designed to tackle the proximity bias. For each chromosome arm, this method subtracts the average LFC value of the hosted unexpressed genes from the original LFC scores.

CCR and MAGeCK can take in input files in FASTQ or BAM format, while all the other methods generally require raw read count files with the only exception of the Geometric method which takes input pre-processed sgRNA depletion LFCs.

Across all the benchmarked methods, CCR and LDO are the only ones working in a completely unsupervised fashion not requiring any additional data other than the CRISPR-Cas9 screen to process. The Geometric method requires the basal expression profiles of the screened model and thus can not be considered a purely unsupervised approach. All remaining algorithms work in a supervised manner, requiring some type of information about the CN status of the genes: MAGeCK, Chronos, AC-Chronos and GAM take in gene-level CN data, while Crispy works with segment-level CN data.

### Overview of the benchmark analysis and employed data

We benchmarked the eight methods described in the previous section by using them to process the two largest to date publicly available Cancer Dependency Map (DepMap) datasets [17, 22]: Project Achilles [45] and Project Score [7, 15, 46] (**Table 2, Methods**). These projects encompass data derived from genome-wide CRISPR-Cas9 viability screens of 1,018 immortalised human cancer cell lines and 18,535 genes (Project Achilles) obtained employing the AVANA sgRNA library [25], and 339 immortalised human cancer cell lines and 18,009 genes (Project Score) screened with the Sanger KY library [6].

**Table 2.**
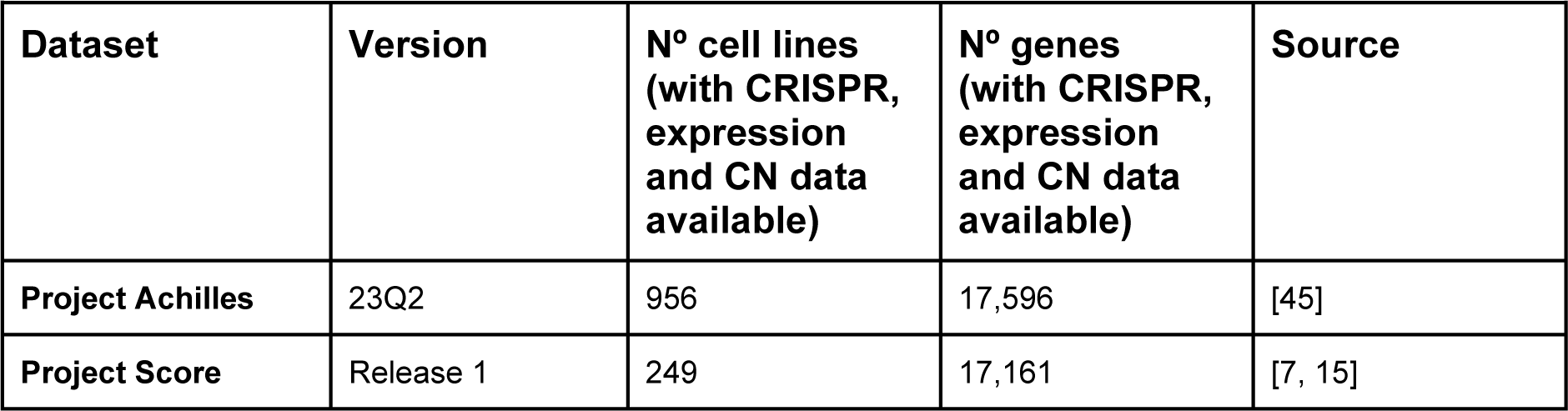
Description of the datasets used to assess the benchmarked methods.

Quality control filtering of these two datasets and removal of cell lines and genes for which CN alteration and basal gene expression data were not available in the DepMap portal (depmap.org) yielded 17,596 genes screened across 956 cell lines for Project Achilles, and 17,161 genes screened across 249 cell lines for Project Score (**Table 2, Methods**).

We tested all the methods with default settings as described in their respective introductory publications and processed the datasets in a multi-screen fashion when the tested method required that. Due to its significant memory and time requirement to process large-scale datasets, we executed MAGeCK on the two tested datasets in batches of optimal size (N = 50), which was determined in a data-driven way (**Fig. S1, Methods**).

We evaluated the ability of each method to reduce CN and proximity biases in the analysed dependency profiles while minimising data distortion, thus preserving screening quality and the ability to detect true positive essential genes. We evaluated the latter aspects by computing quality scores and indicators of true biological signals’ preservation in the screens after they have been processed with the tested methods. Particularly, we computed receiver operating characteristic (ROC) indicators, considering each processed screen as a classifier and highly CN-amplified unexpressed genes, or gold-standard common essential genes as true positive hits, for the two tasks respectively. Finally, we evaluated the ability of the processed screens to detect genomic context-specific dependencies (such as oncogenic addictions) and to unveil significant biomarkers associated with gene dependency, thus preserving data heterogeneity.

### Copy number bias correction comparison

In a reliable, unbiased screen, targeting unexpressed genes with the CRISPR-Cas9 system should not exert any effect on cellular fitness. In a typical (uncorrected) CRISPR-Cas9 screen, CN amplifications generate gene-independent detrimental effects on cellular fitness, leading to highly CN-amplified genes being detected as strongly essential, regardless of their function or expression [30].

In light of this, to evaluate the performances of the tested methods in correcting the CN bias, we focused on cell line specific unexpressed genes (transcripts per million reads mapped, TPM < 1) and grouped them according to their absolute CN value in each individual cell line, using PICNIC processed data [47] from the GDSC database [48]. Then, we compared gene LFCs before and after correction with each of the tested methods across CN bins, assessing the degree to which median values were realigned towards 0. We observed variable results across methods with optimal realignments obtained when processing the two datasets with GAM and CCR (residual difference averaged across CN-bins, ARD = 0.136 and 0.305 respectively, for Project Achilles and ARD = 0.09 and 0.341 respectively, for Project Score). Conversely, we observed reduced yet evident realignments for Chronos, AC-Chronos and Crispy (ARD = 0.829, 0.794 and 0.436 respectively, for Project Achilles and ARD = 0.698, 0.667 and 0.552 respectively, for Project Score, **Fig. 2A and S2A**, **Table S1**, **Methods**).

**Figure 2.**
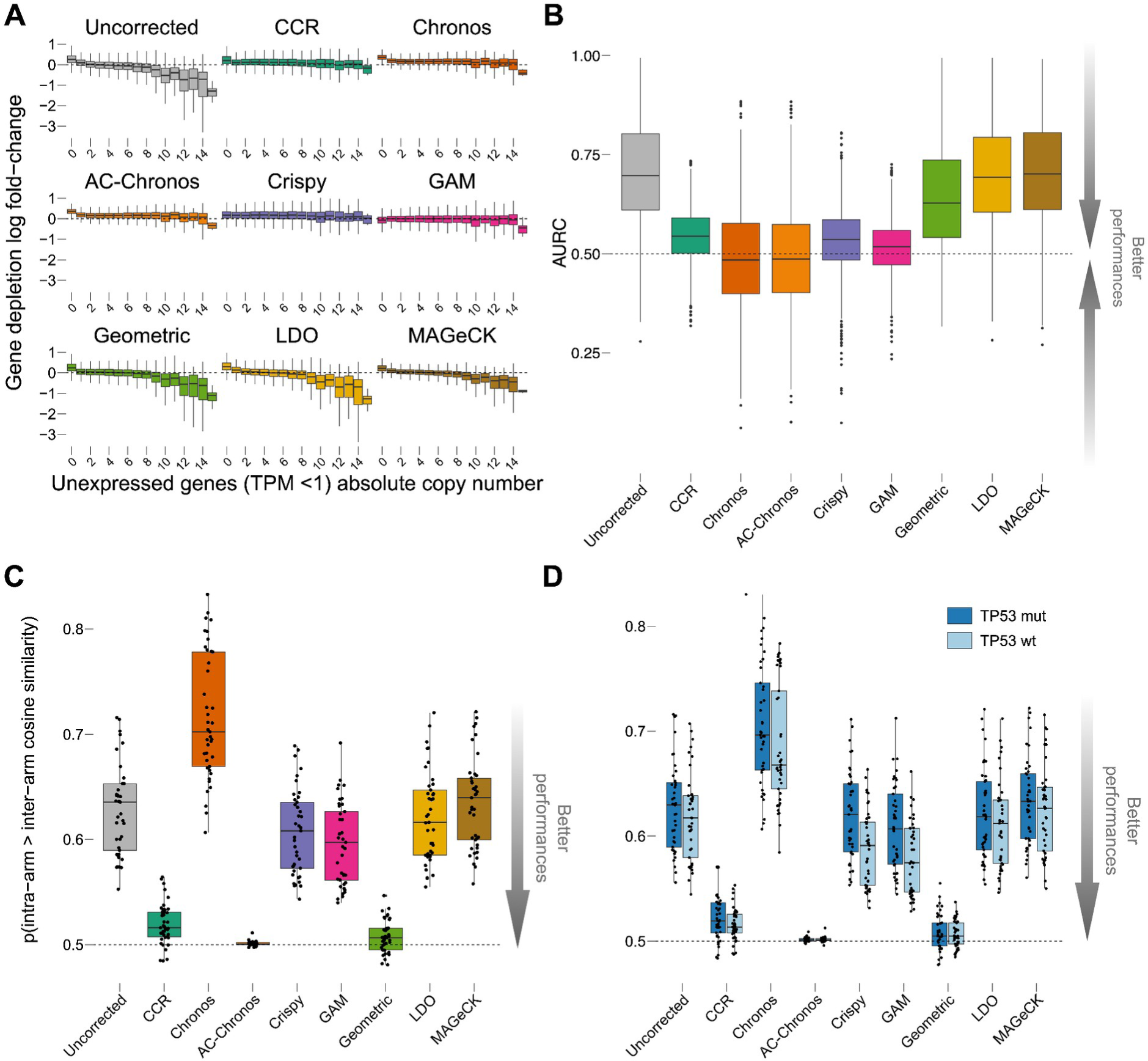
Evaluation of copy number and proximity bias correction for Project Achilles. A. Gene depletion log fold changes (LFCs) of unexpressed genes (TPM < 1) on a per-cell line basis in the Project Achilles dataset grouped according to genes’ absolute copy number value before (uncorrected) and after each method’s correction. B. Screen-wise Area Under the Recall Curve (AURC) of the top 1% amplified unexpressed genes for unprocessed data and across correction methods using the remaining unexpressed genes as outgroup. C. Chromosome arm-level proximity bias quantified by the Brunner-Munzel test statistics for the Project Achilles dataset before (uncorrected) and after each method’s correction. Each point is a chromosome arm. D. Chromosome arm-level proximity bias as in C. but from cell lines split based on the TP53 status. All boxplots represent distributions with central lines indicating median values and lower and upper whiskers encompassing scores within -1.5 and 1.5 times the interquartile range.

Next, for each method and dataset, we considered each processed screen as a rank-based classifier of essential genes (with ranks determined on the basis of their LFCs). We assessed the tendency of such classifiers to predict as essential those genes that were highly CN-amplified (top 1%) but not expressed (TPM < 1) in the screened cell line (median number across cell lines = 7,028), using the remaining unexpressed genes as outgroup and computing areas under the Recall curve (AURC, one for each screen).

For the screens processed with CCR, GAM, Crispy, and especially Chronos and AC-Chronos, we observed AURCs for amplified but non-expressed genes that were closer to what would be expected from a random classifier. This indicated their effectiveness in correcting CN bias.

The screens processed with CCR, Crispy, and GAM exhibited slightly higher average AURC but generally more stable scores across cell lines (mean AURC across cell lines of 0.544, 0.532, and 0.515 for Project Achilles, and 0.543, 0.531, and 0.521 for Project Score, respectively, as shown in **Fig. 2B** and **S2B**, **Methods**). On the other hand, Chronos and AC-Chronos, on average, demonstrated an average AURC more akin to those of a random classifier but with a broader distribution (mean across cell lines of 0.487 and 0.489 respectively, for Project Achilles and 0.512 and 0.516 respectively, for Project Score, as depicted in **Fig. 2B** and **S2B**, and detailed in **Table S1**, **Methods**), indicating a potential inclination towards improved but less consistent performances.

### Proximity bias correction comparison

Several studies have shown that the activity of the Cas9 enzyme may lead to undesired on-target edits in genomic regions proximal to the targeted gene, also known as “proximity bias”, ranging from kilobase-scale deletions to whole chromosome truncation [36–38, 49, 50]. While many efforts have been put forward to mitigate the off-target effects of the Cas9 enzyme, the phenomenon of chromosome loss remains a concern and it is not fully understood. The proximity bias has been recently highlighted by Lazar and colleagues [35], who observed a stronger pan-cancer correlation between dependency profiles of genes located on the same chromosome arm compared to distant genes. The extent to which this phenomenon biases data derived from a genome-wide CRISPR-Cas9 screen is measured by Lazar and colleagues through a generalised Wilcoxon test, also known as Brunner-Munzel test statistic [51], against the null hypothesis that two random numbers (sampled from two different populations) can be equiprobably larger than each other. Particularly, this test quantifies how the pattern of depletion LFCs of genes on the same chromosome arm are significantly more similar than those of genes in different chromosome arms, overall screens in a given dataset.

We employed this method on the Project Achilles and Project Score datasets before and after correction with all tested methods to assess their ability and performances in correcting the proximity bias (**Methods**). Particularly, for each chromosome arm, we derived an intra-arm cosine similarity distribution, comparing the pattern of depletion LFCs of gene pairs in the same arm. Similarly, we derived an inter-arm distribution, comparing patterns of depletion LFCs for gene pairs where the first one was from the chromosome arm under consideration and the second one from another chromosome arm. From these two distributions, we then computed for each chromosome arm a Brunner-Munzel probability of intra-arm similarities being significantly larger than inter-arm ones (BMP).

As expected, we observed probability values larger than 0.5, indicative of larger intra-arm similarity over inter-arm ones, thus the presence of a proximity bias, in Project Achilles and Project Score datasets (unprocessed versions) both (average BMP across chromosome arms = 0.628 and 0.646, respectively for Project Achilles and Project Score, **Fig. 2C and S3A, Tables S2 and S3**).

Across the tested methods, AC-Chronos showed the largest reduction of proximity bias (average BMP across chromosome arms = 0.501 in both datasets, and t-test *p* from contrasting corrected versus uncorrected data = 8.56 x 10^-22^ and 1.11 x 10^-14^, respectively for Project Achilles and Project Score). Unsurprisingly, we observed a large and significant proximity bias reduction when processing both tested datasets with the Geometric method, designed by Lazar and colleagues specifically for this purpose [35] (average BMP across chromosome arms = 0.508 and 0.52, and t-test *p* from contrasting corrected versus uncorrected data = 1.52 x 10^-22^ and 8.69 x 10^-12^, respectively for Project Achilles and Project Score). Processing both datasets with CCR also resulted in a large and significant proximity bias reduction for both datasets (average BMP across chromosome arms = 0.52 and 0.521, and t-test *p* data = 4.69 x 10^-21^ and 7.89 x 10^-13^, respectively for Project Achilles and Project Score). We observed very mild, not significant reductions for the other methods and a significant increase of the proximity bias consistently across both Project Achilles or Project Score when processing them with Chronos only (average BMP across chromosome arms = 0.716 and 0.726, with t-test *p* = 5.59 x 10^-11^ and 2.1 x 10^-5^, respectively, **Fig. 2C and S3A, Tables S2 and S3**).

TP53 expression has been suggested to reduce chromosomal loss in CRISPR-Cas9 editing [52–54] due to its involvement in the DNA repair mechanism. Consistently, we observed an increased intra-arm similarity in screening data derived from cell lines with loss-of-function TP53 mutations compared to TP53 wild-type ones, when considering the uncorrected version of each dataset as well as post-processing them with each tested method (**Fig. 2D and S3B, Table S2 and S3**). Nevertheless, AC-Chronos and Geometric not only yielded the largest reduction of average BMP across chromosome arms but were also able to minimise the difference of this value across TP53 wild-type and TP53 mutant cell lines (average ratio of 9.998 x 10^-1^ and 1.002 respectively, for Project Achilles and AR = 9.992 x 10^-1^ and 1.001 respectively, for Project Score). Notably, we observed a larger residual difference in the proximity bias between TP53 mutant and TP53 wild-type cell lines in the Chronos, Crispy and GAM post-corrected version of Project Achilles and Project Score (AR = 1.029, 1.053, 1.051 respectively, for Project Achilles and 1.043, 1.062, 1.056 respectively, for Project Score).

### Assessment of data distortion

We determined the extent to which the correction methods applied to the Project Achilles and Project Score alter the overall essentiality profiles of the screened cell lines, compromising the ability to derive functional genomics and pharmacogenomics findings from such data. Particularly, we focused on three main features of the processed screens that might be impacted by the correction methods: (i) the detection of common human essential genes among the significantly depleted ones; (ii) the detection of cell line-specific oncogenic addictions, and (iii) the identification of genomic biomarkers of cancer dependencies.

#### Recall of common essential genes and oncogenic addiction relationships

A commonly employed method for evaluating the quality and reliability of a gene essentiality profile obtained from a genome-wide CRISPR-Cas9 screen is to assess the degree to which positive controls, like well-known human essential genes (or core-fitness genes), are identified among the significantly depleted or essential genes [55, 56]. In contrast, negative controls, such as non-essential or non-expressed genes, should not be identified as significantly depleted or essential.

We performed this assessment by means of receiver operating characteristic (ROC) analysis [57] on all the gene essentiality profiles included in Project Achilles and Project Score, before and after processing them with each of the tested correction methods. To this aim, we considered each genome-wide profile of gene LFCs derived from screening a cell line as a rank-based classifier of common essential and non-essential genes [58, 59], with genes’ rank positions determined by their depletion LFC, in increasing order (**Methods**). We then calculated a true positive rate (TPR or Recall), a false positive rate (FPR), and a positive predicted value (Precision) for each rank position. This involved considering all genes from the top of the ranked list down to that specific rank position as predicted essential genes (i.e., predicted positives). We compiled TPR/FPR (or ROC) and precision-recall curves by aggregating values across rank positions.

Finally, we measured the area under the ROC and precision-recall curves (AUROC and AUPRC) and considered these values as final indicators of gene essentiality quality/reliability for both Project Achilles (**Fig. 3AB**) and Project Score (**Fig. S4AB, Methods**).

**Figure 3.**
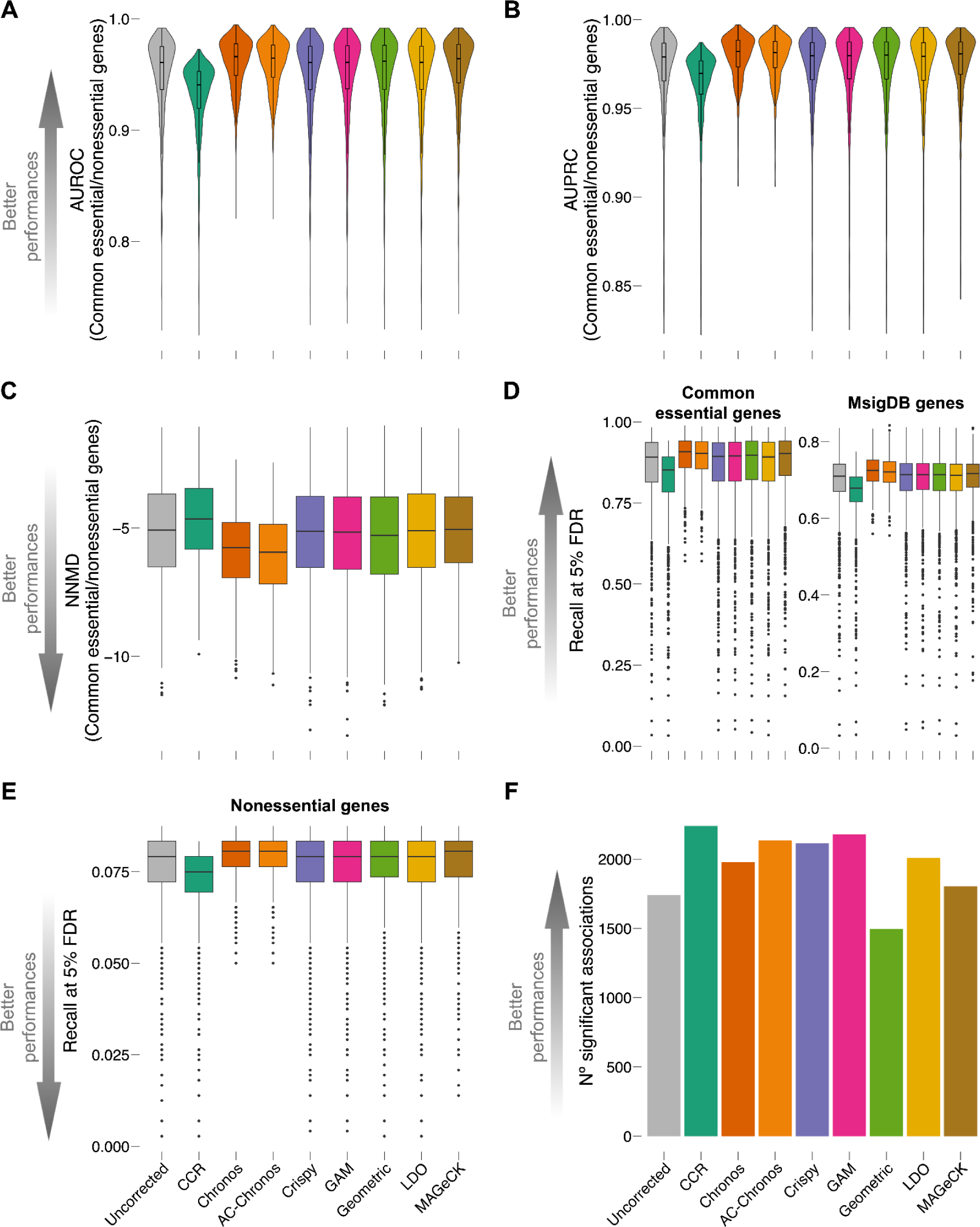
Assessment of data distortion and heterogeneity in Project Achilles. AB. Screen-wise area under the receiver operating characteristic (AUROC) curve (A) and area under the precision-recall curve (AUPRC) (B) obtained when considering all the screens in Project Achilles (uncorrected version and processed with all tested methods) as rank-based classifiers of common-essential/non-essential genes, based on the depletion log fold change (LFC). C. Screen-wise null-normalised median difference (NNMD) between the distribution of LFCs of common essential and non-essential log fold change (LFC) distributions for the screens in Project Achilles (uncorrected version and processed with all tested methods). DE. Screen-wise recall at a 5% false discovery rate was obtained when considering each screen as in ABC but using different gene sets as positive controls (i.e. common essential genes, set of core-fitness genes from MsigDB (E) and non-essential genes (N)) for the Project Achilles dataset (uncorrected version and processed with all tested methods). F. Number of tissue-specific statistically significant (FDR < 5%) Cancer Functional Events (CFEs) / gene-essentiality associations identified in Project Achilles (unprocessed version and across correction methods) via a systematic two-sided *t*-test. This test assesses differential gene essentiality (as measured by the gene depletion log fold-change (LFC)) across two sub-populations of cell lines stratified based on the presence/absence of the considered CFE.

Across all tested methods, Crispy, GAM, Geometric, LDO and MAGeCK substantially preserved the high median AUROC and AUPRC scores observed in the uncorrected version of both Project Achilles (0.949 and 0.972, respectively, **Table S4**) and Project Score (0.936 and 0.969, respectively, **Table S5**). Chronos and AC-Chronos slightly improved these scores (0.961 and 0.96 respectively, for AUROC in Project Achilles; 0.954 and 0.952 respectively, for AUROC in Project Score; 0.98 and 0.979 respectively, for AUPRC in Project Achilles; 0.978 and 0.977 respectively, for AUPRC in Project Score) consistently with these methods analysing multiple screens in a joint fashion, which boosts the signals of genes that tend to be essential across cell lines (thus of common essential genes), improving the quality of screens exhibiting a lower phenotype penetrance. In contrast, CCR slightly reduced AUROC and AUPRC for both Project Achilles (0.93 and 0.963, respectively) and Project Score (0.92 and 0.964, respectively), consistently with its unsupervised approach that smooths depletion signals overall.

An alternative method for assessing the quality and reliability of genome-wide essentiality profile derived from a CRISPR-Cas9 screen consists of evaluating how well separated the distributions of depletion LFCs of common essential and non-essential genes are. A measure of this separation is the null-normalised mean difference (NNMD) [45, 46] defined as the difference between the median of the common essential and non-essential genes’ LFC distributions divided by the median absolute standard deviation of the non-essential genes’ LFC distribution, which yields large negative values for high-quality data (**Methods**). We observed large negative median NNMD values for the uncorrected versions of Project Achilles and Project Scores (-5.19 and -5.04, respectively), which were largely preserved following correction with the majority of the tested methods, mildly improved by Chronos (-5.93 and -5.74, respectively) and especially AC-Chronos (-6.09 and -5.91, respectively), and slightly worsened by CCR (-4.7 and -4.91, respectively, **Fig. 3C and S4C, Table S4 and S5**).

These results were confirmed when extending our analysis to the Recall at a fixed level of 5% false discovery rate (FDR) and including additional sets of prior known essential genes (from Pacini *et al*. [46] and the molecular signature database (MsigDB) [60], assembled as detailed in Iorio *et al*. [34]) (**Fig. 3D and S4D, Tables S6 and S7, Methods**).

Finally, we assessed the extent to which the tested correction methods impacted the detection of false positive essential genes. To this aim, we computed the Recall of non-essential genes (from [58]) at a fixed level of 5% FDR, across individual screens before and after the correction of both tested datasets with all considered methods. Also in this case, values were largely conserved across all tested methods and proximal to zero, i.e. median ∼0.08 for both datasets, before and after correction with all tested methods. Of interest is the comparison between CCR and LDO as they stand as the only unsupervised single-screen methods. This becomes paramount when researchers are confronted with constraints such as a limited availability of CRISPR screens and an absence of additional omics data (e.g CN data). Notably, CCR demonstrates better ability over LDO in addressing CN and proximity biases associated with CRISPR-Cas9 screening data. Conversely, LDO exhibits a lower impact on data quality upon correction as elucidated by higher recall of common essential genes, while CCR shows a lower recall of nonessential genes, effectively trading sensitivity for specificity.

Next, we examined the ability of each screen to detect known selective dependencies across cell lines and how this was impacted by the assessed correction methods. The constitutive function of oncogenes harbouring gain-of-function mutations tends to become addictive to the cancer model harbouring these alterations. As such, mutated oncogenes tend to be detected as selectively essential when genomically altered in cancer cells. Based on this, we implemented an approach similar to that pursued in our previous work [46] and sought to assemble a set of positive/negative selective dependencies, considering a set of oncogenes from OncoKB [61] (338 in total, of which 329 was screened in Project Achilles, and 327 in Project Score, **Methods**). First, we scaled genome-wide LFC profiles for better cross-screen comparability (i.e. median LFC score of -1 and 0, respectively for common essential and non-essential genes across cell lines) as done in previous works [46, 56]. We then determined the status of these oncogenes in each screened cell line focusing on each of the two tested datasets in turn (using their multi-omic characterisation from the DepMap portal), collating cell-line specific mutated oncogenes in a collective set of positive controls (n = 7,861 and 2,210, respectively for Project Achilles and Project Score) and cell-line specific wild-type and unexpressed (TPM < 1) in a collective set of negative controls (n = 81,577 and 19,456, for Project Achilles and Project Score respectively). Next, we pooled all oncogenes’ LFCs across screens in a unique list of 290,624 and 72,957 entries, respectively for Project Achilles and Project Score, and considered it as a rank-based classifier of selective dependencies, with gene rank positions determined on the basis of the gene’s scaled LFC, using the assembled list collective positive/negative controls. Finally, we assessed the performances of such a classifier by means of ROC analysis, computing AUROCs for both datasets before and after processing them with each tested method.

We observed slightly better than random performances for both Project Achilles and Project Score (uncorrected versions, AUROC = 0.649 and 0.641, respectively, **Figure S5A** and **S5B**) which were mostly preserved by the Geometric, LDO and MAGeCK methods, with MAGeCK resulting as the top-performing method for Project Achilles (AUROC = 0.655), and LDO and Geometric for Project Score (AUROC = 0.647) (**Tables S4 and S5**).

#### Identification of biomarkers

The identification of context-specific genetic dependencies is one of the key aims of cancer pharmacogenomics and is pivotal for the development of future therapies with associated predictive markers. These can be identified in CRISPR-Cas9 screening datasets by focusing on genes eliciting a strong reduction of viability in a subset of cell lines upon CRISPR-Cas9 knockout. These context-specific dependency genes often show a statistically significant differential dependency when contrasting cell lines based on the presence/absence of a given molecular feature (such as a gene mutation or CN amplification). Such features can then be used as the basis for the development of future therapeutic biomarkers, might provide mechanistic insights on the associated genetic dependency and unveil involved biological pathways [7, 28, 46].

We investigated the extent to which tissue-specific biomarkers associated with cancer dependencies can be identified in the unprocessed datasets and following the application of each benchmarked method (**Methods**).

To achieve this, we considered a set of 1,073 tissue-specific clinically relevant cancer functional events (CFEs), including somatic mutations in well-established cancer driver genes, genomic segments that are recurrently CN-altered in cancer, and hypermethylated gene promoters as identified in our previous work [48]. We considered 25 different tissues. For each possible combination of CFE, tissue, processing method, and dataset, we performed a Student’s *t-test* to evaluate the differential essentiality of a strongly selective dependency (SSD) gene (**Methods**). This involved comparing the gene’s depletion LFCs across two sub-populations of screened cell lines from the specific tissue and dataset (uncorrected or processed with the chosen method), stratified based on the presence or absence of the considered CFE. We tested a total of 3,553,680 and 2,183,328 CFE/SSD pairs with suitable sample sizes (i.e. at least 3 cell lines in the contrasted subpopulations, **Fig. 3F** and **S4F**).

Most of the tested correction methods increased the number of significant (< 5% FDR) CFE/SSD associations compared to those detectable in the unprocessed versions of both datasets. CCR processed Project Achilles dataset unveiled the highest number of significant associations (n = 2,240) compared to the other methods, while Chronos, AC-Chronos and Geometric processing performed the best for Project Score by a large margin (n = 645, 637 and 620, respectively), followed by CCR (n = 493, **Table S8**).

## Discussion

Genome-wide CRISPR-Cas9 screens have increasingly become essential tools in functional genomics and early drug discovery, owing to their potential to unveil synthetic lethal gene pairs or genes responsible for drug resistance/sensitivity. However, correctly interpreting and analysing data from such screens is hampered by false positive hits arising from technology-specific biases. These relate to the mode of action of the Cas9 enzyme, which generates gene-independent viability reduction responses when targeting copy number amplified genomic regions [30] or are due to whole chromosome harm truncations [35] (causing genes in proximal positions to elicit similar effects on cellular fitness upon CRISPR-Cas9 targeting regardless of their function or expression).

Many algorithms have been proposed to correct such biases *in silico*. Here, we performed a benchmark analysis assessing the most recent eight of these [24, 33–35, 39–42, 62] and assessed their bias correction performances. We also estimated their impact on data quality under different use cases. Although there have been previously published studies benchmarking common algorithms for the analysis of pooled CRISPR screens [63], our analysis is the first one thoroughly benchmarking recently published methods specifically designed to address biases in CRISPR-Cas9 screening data associated with established [30] or recently reported [34, 35] structural features of the targeted genomic regions.

Among the tested methods, we found that Chronos showed lower effectiveness in proximity bias correction (**Fig. 4A**). CCR, GAM, Crispy and especially Chronos and AC-Chronos obtained the best performances in addressing the CN bias. On the other hand, we observed no clear distinction in performance between supervised (i.e. requiring CN or other kind of omics data) and unsupervised methods. A possible explanation is that CRISPR-Cas9 screening data are affected by multiple source of biases and supervised methods are designed to address only one (i.e. CN bias) or two (i.e. CN and proximity biases) of them, missing others such as off-target effects or differences in cellular responses. Of note, CCR, not requiring any additional information beyond the (individual) CRISPR-Cas9 screen to process, shows good performances in correcting both the CN- and proximity-bias, while Geometric, needing basal expression profiles of the screened models and genomic target coordinates of the employed sgRNA library, efficiently removes the proximity bias but tends to under-correct the CN bias. However, we observed that AC-Chronos, the new pipeline developed on top of the original Chronos, is the best at addressing both the CN and proximity biases.

**Figure 4.**
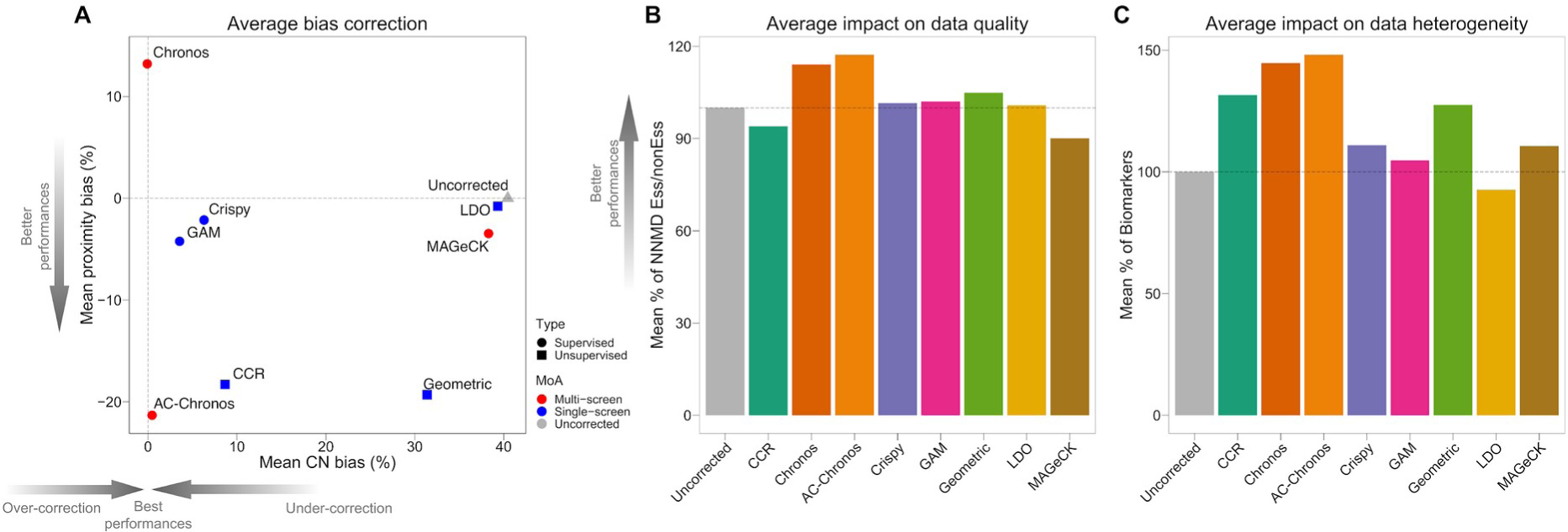
Summary of tested methods’ ability to correct for structural biases associated with CRISPR-Cas9 screens and their impact on data quality and heterogeneity. A. Tested methods’ performances (individual points) in correcting copy number (CN) bias (x-axis) and proximity bias (y-axis). Coordinate on the y-axis indicate the mean reduction of proximity bias with respect to uncorrected data averaged across datasets. Coordinates on the x-axis indicate the mean area under the recall curve (AURC) considering the correction as percentage change with respect to the uncorrected version, and averaged across datasets. B. Tested methods’ performances (individual points) in addressing the impact on data quality: mean null-normalised median difference (NNMD) (y-axis) of common essential and non-essential genes expressed as percentage change with respect to the uncorrected version, and averaged across datasets. C. Summary of tested method’s performances (individual points) in addressing the impact on data heterogeneity: number of tissue-specific statistically significant (FDR < 5%) Cancer Functional Events (CFEs) / gene-essentiality associations (y-axis) expressed as percentage change with respect to the uncorrected version, and averaged across datasets. For all plots, each dot is a method: the shape corresponds to the type of additional data required for correction, supervised (i.e. requiring CN scores) or unsupervised, and the colour corresponds to the mode of action, single- or multi-screen.

Further, we estimated the distortion introduced by each method’s correction on the gene essentiality profiles derived from each processed screen, considering them as rank-based classifiers of pre-defined sets of core-fitness-essential/non-essential genes (with gene rank positions determined by the extent of viability reduction observed upon CRISPR-Cas9 targeting) (**Fig. 4B**). Finally, we assessed the extent to which each correction method preserved the ability of the processed datasets to unveil synthetic lethalities and markers of gene essentiality (**Fig. 4C**).

Results confirmed that, by modelling the essentiality signal of each gene by borrowing information across screens in the same dataset, AC-Chronos and Chronos yield corrected screens that are generally better predictors of core-fitness essential genes. In particular, AC-Chronos is the best method for the correction of both types of structural biases and it also yields more stable correction outcomes compared to the other methods. On the other hand, the correction performed by CCR comes at the expense of the gene essentiality signals which are slightly smoothed in the individual screens leading to weaker performances in classifying essential/non-essential genes. In addition, AC-Chronos and Chronos seem to also preserve individual screens’ heterogeneity, allowing for the identification of a larger number of biomarker/gene-essentiality associations compared to other methods. They are also better able to improve data quality overall and they are best suited for modelling complex experimental settings (for example, accounting for gene essentiality data at multiple time points or integrating multiple screens performed with different sgRNA libraries).

## Conclusions

Our benchmark unveiled the strengths and weaknesses of each method across the main factors that may affect the analysis and interpretation of data derived from CRISPR-Cas9 screens. Based on our results AC-Chronos should be the primary choice for the correction of CN and proximity biases as well as for its ability to minimise the impact on data quality, when processing multiple screens with available CN-information jointly. AC-Chronos also allows the identification of a larger number of total clinically relevant biomarker/dependency associations. Nonetheless, CCR offers very good performances for the correction of CN and proximity biases and its ability to work on a single-screen basis without any additional omic data makes it the method of choice in case of limited data availability.

On the other hand, the corrected datasets yielded by AC-Chronos and Chronos, which borrows information across screens, had a lower impact on the data quality and, thus, we were better able to recapitulate priori known common essential and non-essential genes.

Overall, there is no one-size-fits-all method that outperforms the others across the analyses we carried out, rather it is up to the researchers to carefully pick the method best suited for their needs according to their use case. Finally, our analyses can be applied to forthcoming and additional CRISPR-Cas9 datasets, enabling a more extensive coverage of each method’s correction performances.

## Methods

### Pre-processing of cancer dependency datasets

We focused on the two largest genome-wide CRISPR-Cas9 screen datasets to date: Project Score (release 1), screened using the KY library [64, 65], and Project Achilles release 23Q2, screened using the AVANA library [66]. We downloaded the raw read count versions of both datasets from the DepMap portal (https://depmap.org/portal/download/all/). These datasets were already pre-processed with the DepMap pipeline up to the computation of raw read counts (README.txt, https://depmap.org/portal/download/all/), yielding 1018 cell lines and 18,535 for the Project Achilles dataset, and 339 cell lines and 18,009 genes for the Project Score dataset. We then applied the following procedure:

- Single-guide RNAs (sgRNAs) and technical replicates that did not pass the quality control were removed based on the quality report files available on the DepMap web portal (i.e. [Lib]GuideMap.csv file for the sgRNAs quality, where [Lib] stands for the library used to generate a given dataset (“Avana” or “KY”); AchillesSequenceQCReport.csv file for the quality of the technical replicates of both datasets). Low-quality sgRNAs were defined as those having multiple alignments to the same gene or multiple genes, no alignments, only intergenic alignments, or being the only passing guide targeting a gene. Low-quality replicates were defined as those having less than 185 mean read counts per guide, a Pearson coefficient < 0.41 with at least one other replicate when looking at genes with the highest variance in gene effect across cell lines, and a Null-normalised median difference (NNMD) from the log fold change (LFC) > -1.25. NNMD is computed as follows:

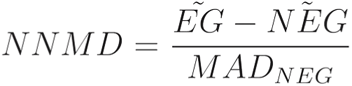

where 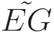 and 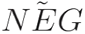 are the median LFC values of the essential and non-essential reference gene sets, and MAD_NEG_ is the median absolute deviation of the non-essential reference genes as computed in [46]. This yielded 971 cell lines and 17,787 for the Project Achilles dataset, and 317 cell lines and 17,349 genes for the Project Score dataset.
- For raw read counts, NAs were replaced with 0.

Considering each method has different input requirements, we derived the corrected datasets as follows: we ran MAGeCK MLE directly on the raw read counts as it required to implement a design matrix contrasting for each cell line the technical replicates post-library transduction against the reference plasmid DNA counts. CRISPRcleanR and Crispy were run on the sgRNA-level raw LFCs, applying the following steps:

- LFCs were computed by scaling the raw read counts post-library transduction by the reference plasmid DNA counts for each replicate.
- LFC scores per guide were averaged across high-quality replicates to obtain sgRNA-level LFCs for each cell line.

Geometric, LDO and GAM methods were applied to gene-level LFCs and the datasets were further processed as follows:

- Raw gene-level LFCs were obtained by taking the median value among the same gene-targeting sgRNAs per screened cell line.

Chronos, instead, needs to be trained on the dependency data to optimise the parameters of its mechanistic model of cell population dynamics. This includes the removal of clonal outgrowths and the inference of uncorrected gene effects [39]. To avoid retraining the model as the results were already available, we downloaded the inferred gene effects from the DepMap portal (ScreenGeneEffectUncorrected.csv, https://depmap.org/portal/download/all/). Then we applied for each dataset the Chronos correction strategy for copy number bias.

Finally, we intersected the corrected dependency datasets considering only cell lines and genes in common for the quantification of biases and data distortion analyses. This yielded 956 cell lines and 17,596 genes for Project Achilles, and 249 cell lines and 17,161 genes for Project Score.

### Arm-corrected Chronos

To correct the proximity bias post-Chronos processing, the median gene effect of each chromosome arm is aligned to be the same across all screens. This involves finding the median gene effect per screen *m_s_* for all genes *g* in a given chromosome arm within a cell line. The difference between *m_s_* and the median med(*m_s_*) across all cell lines is then subtracted from the gene effect of all genes on the arm *e_g_*, yielding the corrected gene effect for that arm as follows:

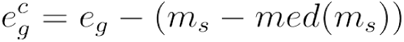

This correction is applied to all chromosome arms with > 5 genes.

### In-house implementation of the Geometric method

As the code for the Geometric method has not been made publicly available yet, we implemented our in-house version of this method following the description provided in Lazar *et al*. [35] and considered as unexpressed those genes with a transcript per million (TPM) < 1 for a given cell line in the DepMap release 23Q2 [45], using basal RNAseq bulk profiles of the same cell line available on the DepMap portal (https://depmap.org/), preprocessed following the DepMap Omics processing pipeline available at: https://github.com/broadinstitute/depmap_omics.

### Batch-wise execution of MAGeCK MLE

Unlike other tested methods, MAGeCK MLE’s execution on the entire Project Achilles and Score datasets required prohibitive amounts of memory and time, making it impractical to execute the correction in a single run. For example, when tested on the entire Project Achilles dataset on the institute’s in-house high-performance computing cluster, allocating 300 GB of memory and the maximum computing time available (i.e. two weeks) the job still failed due to an out-of-memory issue.

To take full advantage of MAGeCK MLE’s capabilities to process and correct CRISPR-Cas9 knock-out screens in a pooled fashion, we devised a practical solution by running the method in batches of screens. First, we explored the impact of this batching strategy on result stability and accuracy as follows: we randomly split the dataset (i.e. Project Achilles or Project Score) into batches of different sizes (i.e. 5, 10, 20 and 50 cell lines) at a fixed seed and combined the corrected batches back into a final dataset. We then computed the pointwise difference between MAGeCK beta scores obtained using batches of different sizes (**Fig. S1AB**). This is defined as the difference between the same cell-gene coordinates across datasets of different batch sizes. We observed that the bulk of the distribution was very narrow and centred at around 0 with the differences between MAGeCK beta scores shrinking across datasets as we increased the batch size. In particular, when binning these pointwise differences between the respective datasets processed in batches of 20 and 50 cell lines, more than 99.9% were less than 0.1, considering beta scores have a range approximately between -7 and 9 for the datasets under consideration (**Fig. S1CD**). We then focused on a batch size of 50 as it was the largest amount of cell lines possible to process without out-of-memory issues.

Finally, to test MAGeCK robustness we derived 100 corrected datasets, processed in batches of 50 cell lines, using different seed values each time and computed the pointwise standard deviation (**Fig. S1EF**). This is defined as taking each time the same cell-gene coordinates across seeded datasets and computing the standard deviation on the resulting vector. The vast majority of data points show great stability and very low standard deviation. Thus, these results motivated us to process the datasets in batches of 50 cell lines for MAGeCK MLE.

### Quantification of copy number bias

For each dataset, we assessed each method’s capability to correct for copy number (CN) bias by computing the area under the recall curve (AURC) of the amplified non-expressed gene sets. These genes were defined as those in the top 1% highest CN score which are not expressed (TPM < 1) for a given cell line. Remaining non expressed genes were used as outgroup. The AURC is independent of a fixed LFC threshold. For each rank position *k*, we derived the corresponding threshold-specific recall. Finally, we applied the trapz function from the pracma R package (v2.4.2) on the set of recall scores to obtain the AURC.

In addition, we grouped gene-level LFCs by their PICNIC absolute CN score [47] from the GDSC resource [48]. Then, for each method, we computed the average residual difference (ARD) across CN-binned LFCs as follows:

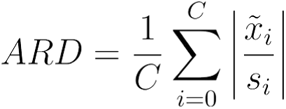

where 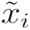 and *s_i_* are the median score and the standard deviation derived from the LFC distribution binned for *i^th^* absolute CN value of a given method, respectively. *C* is the maximum absolute CN value. This metric indicates the goodness of a method in centring the LFC distributions towards 0.

### Quantification of proximity bias

For each method and dataset, after correction, we computed the pairwise cosine similarity between gene targets and ordered them by chromosome position along the human genome. The results were then quantile normalised in order to have a mean of 0 and a standard deviation of 0.2, following the same procedure by [35]. To test the proximity bias according to TP53 status, we first split the datasets in TP53 wild-type and mutated cell lines and repeated the aforementioned pipeline on each block.

Next, we utilised the Brunner-Munzel test statistic [51] to assess the presence of proximity bias on a genome-wide scale. For each chromosome arm, we compared the intra-arm cosine similarity distribution, where both genes belong to the same arm, and the inter-arm distribution, where only one of the two genes belongs to the arm under consideration while the other belongs to any other chromosome arm.

### Recall of common essential genes

We considered two sets of a priori known common essential and non-essential genes as positive and negative classes to estimate the impact of each method’s correction on the data quality. These sets were downloaded from the DepMap portal (https://depmap.org/portal/download/all/): files AchillesCommonEssentialControls.csv, resulting from the intersection of the Hart and Blomen essential genes [5, 59], and AchillesNonessentialControls.csv, containing the Hart reference non-essentials [58]. For each dataset, algorithm and cell line, we computed the area under the receiver operating characteristic (AUROC) and the area under the precision-recall curve (AUPRC) using the ccr.ROC_Curve and ccr.PrRc_Curve functions in the CRISPRcleanR R package (https://github.com/francescojm/CRISPRcleanR)[34, 62]. Final sets of positive and negative controls were determined by intersecting the common essential/non-essential gene sets with the genes screened in the dataset under consideration (either Project Achilles or Project Score) across all algorithms. In addition, we calculated the cell-wise NNMD between the two gene sets as the difference in the median LFC of positive and negative controls, normalised by the median absolute deviation of the negative controls.

### Dependency biomarkers’ analysis

We downloaded a comprehensive table of annotated cancer genetic variants in cancer from the OncoKB database (https://www.oncokb.org/cancerGenes, 12/21/2023 as last update). To ensure the relevance of the data, we focused on genes marked as having gain-of-function or switch-of-function alterations (i.e. annotated genes with a clear oncogenic role). First, we scaled the genome-wide essentiality profiles for each method and dataset tested so that the median LFC scores of common essential and non-essential genes were -1 and 0, respectively, across cell lines, allowing cross-screen comparability.

For each specific oncogene, we utilised somatic mutation and fusion calls from the DepMap portal release 23Q2: oncogenes found mutated in a given cell line were deemed as positive controls, while those that were found wild-type and not expressed (TPM < 1) were considered as negative controls. In addition, only oncogenes with at least one positive and one negative control were retained. Finally, for each method and dataset we calculated the AUROC curve on the pooled oncogene LFC scores using the ccr.ROC_Curve function in the CRISPRcleanR R package (https://github.com/francescojm/CRISPRcleanR)[34, 62] to estimate each method’s ability to recall oncogenic addictions.

### Recall of predefined gene sets of essential/non-essential genes

We considered three predefined sets of genes: common essential and non-essential genes derived from the DepMap portal, and additional essential genes derived from the Molecular Signature Database (MsigDB) (https://www.gsea-msigdb.org/gsea/msigdb) [60]. To generate MsigDB sets of prior essential genes we downloaded gene sets from [46], originally derived from MSigDB (v7.2). The gene sets used from KEGG were KEGG_DNA_REPLICATION, KEGG_RNA_POLYMERASE, KEGG_SPLICEOSOME, KEGG_RIBOSOME and KEGG_PROTEASOME. For the histone gene set, we combined two Reactome gene sets, namely REACTOME_HATS_ACETYLATE_HISTONES and REACTOME_HDACS_DEACETYLATE_HISTONES as well as the curated histones gene set from [7].

For these gene sets, we computed the recall at 5% false discovery rate (FDR) across methods and datasets. Considering an essentiality profile, we ranked the LFC scores of the reference common essential (E) and non-essential (N) genes in increasing order. For each ranked position *k*, a set of predicted essential genes is found:

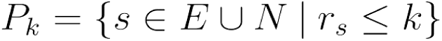

where *r_s_* indicates the rank position of *s*, and the corresponding positive predictive value (PPV) is computed as follows:

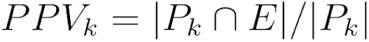

We then determined the lowest threshold of LFC (*LFC\**) in rank position *k\** with *PPV_k_* ≥ 0.95. This is equivalent to an FDR ≤ 0.05. The *LFC\** threshold corresponds to *r_s_* = *k\**. All genes with an LFC score ≤ *LFC\** were deemed essential at 5% FDR. Given one of the three gene sets *G* (i.e. common essential, MsigDB essential or non-essential), the recall at 5% FDR is defined as:

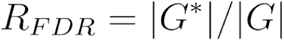

where *G\** is the subset of genes in G below the *LFC\** threshold.

### Systematic association between cancer functional events and gene dependencies

For each method, dataset, CFE and lineage, we systematically performed a two-sided unpaired *t*-test to assess the differential essentiality of each strongly selective dependency (SSD) against the status (presence/absence) of predefined cancer functional events (i.e. somatic mutations, recurrently aberrant copy number segments and hypermethylated sites) [48]. These SSDs were identified using a likelihood ratio test as described in [28, 46]. The null hypothesis assumed equal means in the compared populations, while the alternative hypothesis suggested an association between the tested CFE/gene-dependency pair. To account for multiple hypothesis testing, we corrected the obtained p-values using the Benjamini–Hochberg method. The associations with an FDR of less than 5% were deemed significant.

## Availability of data and materials

The code to reproduce results and figures is available at: https://github.com/AleVin1995/Structural_bias_benchmark. The code is provided under the GPL-3.0 license. The datasets analysed in this manuscript are publicly available on the DepMap website at: https://depmap.org/portal/download/all/. They include Project Achilles 23Q2 and Project Score release 1, as well as somatic mutation, copy number, gene fusion, gene expression and all the metadata related to CRISPR-Cas9 screens (i.e. screen quality control, guide to gene mapping and sequence to screen mapping). In addition, the OncoKB cancer driver gene list can be found at: https://www.oncokb.org/cancerGenes, and the PICNIC absolute copy number scores at: https://cog.sanger.ac.uk/cmp/download/cnv_20191101.zip.

## Acknowledgements

We thank Ludovica Proietti and Gianluca Vozza for their critical review of our manuscript and for their constructive feedback. Panels A and C of Figure 1 were created with <u>BioRender.com</u>.

## Author information

### Authors and Affiliations

Computational Biology Centre, Human Technopole, Viale Rita Levi-Montalcini, 1, 20157 Milano, Italy.

Alessandro Vinceti, Raffaele Iannuzzi, Lucia Trastulla, Francesco Iorio.

Cancer Dependency Map Analytics, Wellcome Sanger Institute, Wellcome Genome Campus, Hinxton, Cambridge CB10 1SA, UK.

Francesco Iorio.

Cancer Data Science, Broad Institute of Harvard and MIT, MA, USA

Isabella Boyle, Catarina D. Campbell, Joshua Dempster

Cancer Dependency Map, Broad Institute of Harvard and MIT, MA, USA

Francisca Vazquez

### Contributions

Conceptualisation, A.V. and F.I.; methodology, A.V. and F.I.; software, A.V. and I.B.; formal analysis, A.V., R.I., and L.T.; investigation, A.V.; data curation, A.V. and I.B.; writing – original draft, A.V. and F.I.; writing – review & editing, A.V., R.I., L.T., I.B., C.D.C., F.V., J.D., and F.I.; visualisation, A.V., R.I., L.T., and F.I.; co-supervision, C.D.C., F.V., J.D.; supervision, F.I.

### Corresponding author

Correspondence to Francesco Iorio.

## Ethics declarations

### Ethics approval and consent to participate

Not applicable.

### Consent for publication

Not applicable.

