## Supplementary Material for "A benchmark of computational methods for correcting biases of established and unknown origin in CRISPR-Cas9 screening data"

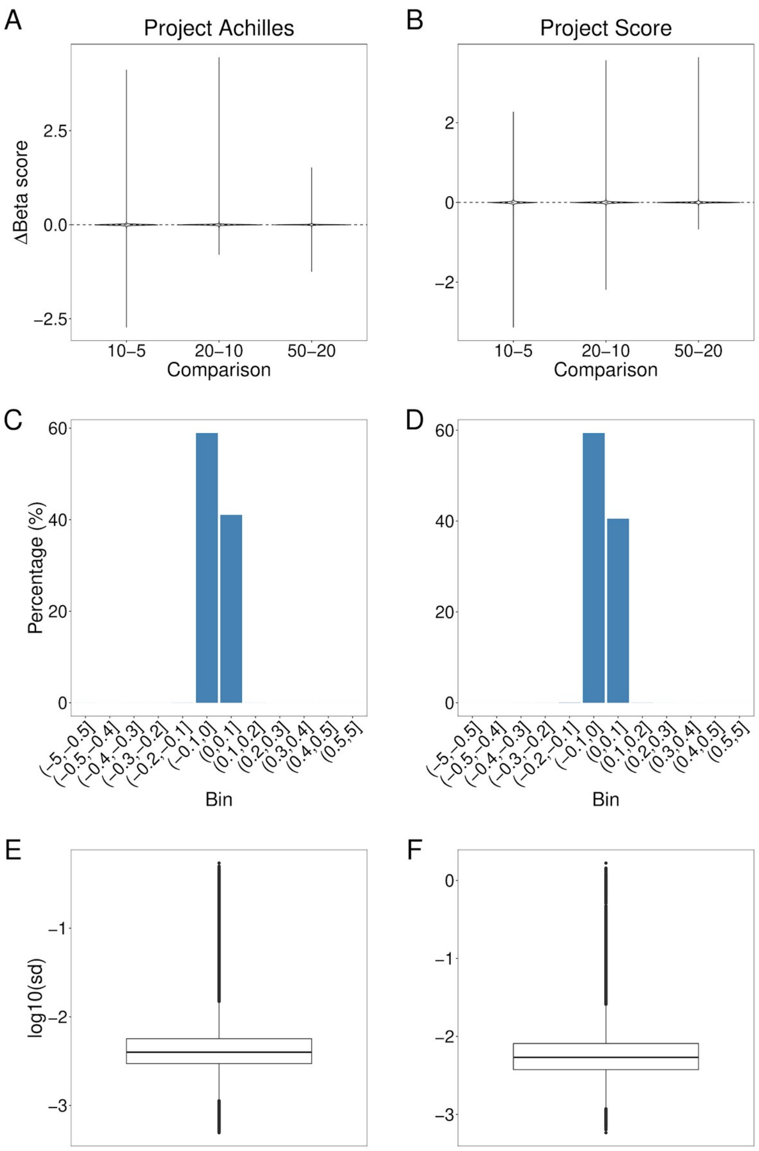


**Figure S1 - Impact of batching on MAGeCK's correction outcome. AB. Pointwise difference between MAGeCK-corrected Project Achilles (A) and Project Score (B) datasets processed in different batch sizes (i.e. 5, 10, 20 and 50 cell lines). CD. Binning of pointwise differences for Project Achilles (C) and Project Score (D) datasets, processed in batches of 20 and 50 cell lines. EF. Pointwise standard deviation across 100 MAGeCK-corrected Project Achilles (E) and Project Score (F) datasets, processed in batches of 50 cell lines, using different seed values.**

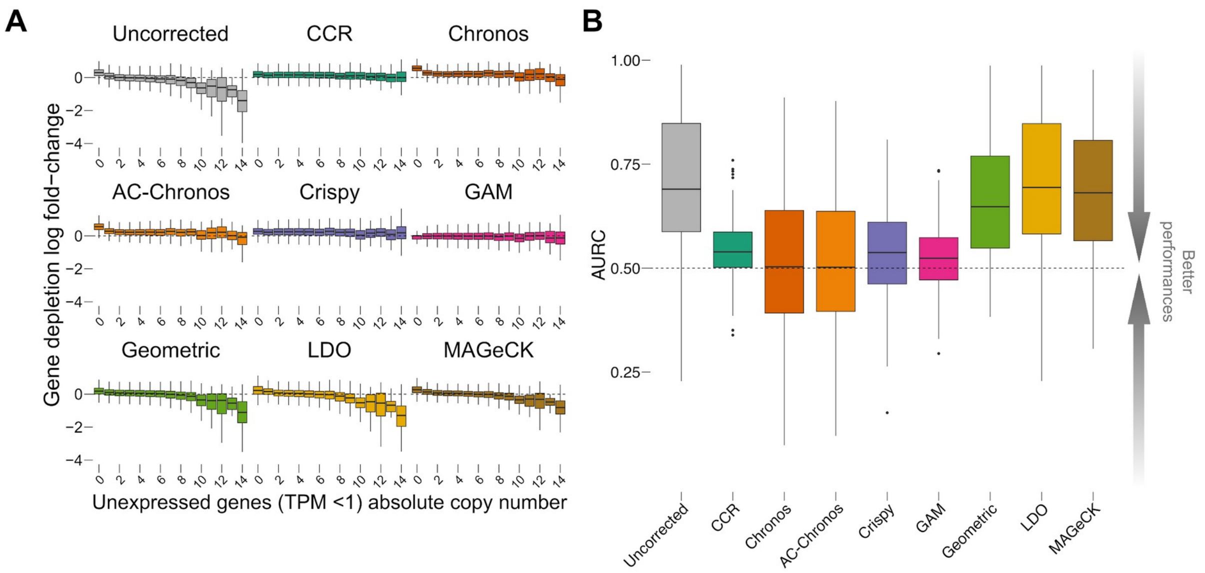


**Figure S2 - Correction of copy number bias for unexpressed genes in Project Score. A. Gene-level log fold-change (LFC) values of unexpressed genes (TPM < 1) on a per-cell line basis in the Project Score dataset grouped according to their absolute copy number value (using PICNIC data) before and after each method correction. B. For each method, we computed the cell-wise Area Under the Recall Curve (AURC) of the top 1% amplified unexpressed genes using the remaining unexpressed genes as outgroup.**

| Method | Project Achilles | | Project Score | |
|---|---|---|---|---|
| | ARD | AURC (Top 1% amplified unexpressed genes) | ARD | AURC (Top 1% amplified unexpressed genes) |
| Uncorrected | 0.66±0.828 | 0.701±0.13 | 0.508±0.454 | 0.703±0.16 |
| CCR | 0.305±0188 | 0.544±0.0711 | 0.341±0.19 | 0.543±0.0716 |
| Chronos | 0.829±0.498 | 0.487±0.137 | 0.698±0.473 | **0.512±0.174** |
| AC-Chronos | 0.794±0.455 | **0.489±0.132** | 0.667±0.451 | 0.516±0.167 |
| Crispy | 0.436±0.213 | 0.532±0.0915 | 0.552±0.237 | 0.531±0.108 |
| GAM | **0.136±0.318** | 0.515±0.07 | **0.0908±0.107** | 0.521±0.081 |
| Geometric | 0.507±0.666 | 0.645±0.126 | 0.381±0.333 | 0.669±0.14 |
| LDO | 0.628±0.794 | 0.695±0.128 | 0.462±0.401 | 0.698±0.159 |
| MAGeCK | 0.673±0.868 | 0.703±0.13 | 0.488±0.408 | 0.68±0.153 |

**Table S1 - Summary of average residual divergence (ARD) and area under the recall curve (AURC) of copy number bias correction (mean ± standard deviation) for unexpressed genes across methods in Project Achilles and Project Score. Bold is the best outcome across methods.**

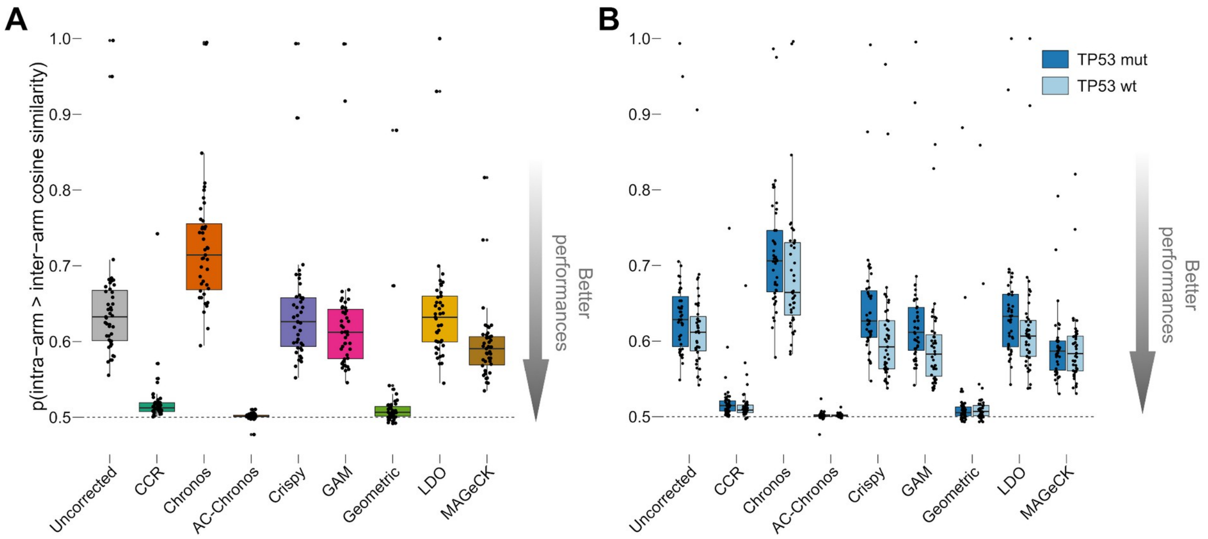


**Figure S3 - Correction of proximity bias in Project Score. A. Chromosome arm-level proximity bias quantified by the Brunner-Munzel test statistics for the Project Score dataset before (uncorrected) and after each method's correction. Each point is a chromosome arm. B. Chromosome arm-level proximity bias as in A. but from cell lines split based on the TP53 status. All boxplots represent distributions with central lines indicating median values and lower and upper whiskers encompassing scores within -1.5 and 1.5 times the interquartile range.**

| Method | GW | P-val | TP53 WT | P-val | TP53 LOF | P-val | AR |
|---|---|---|---|---|---|---|---|
| Uncorrected | 0.628±0.0424 | NA | 0.627±0.0423 | NA | 0.618±0.0419 | NA | 1.015 |
| CCR | 0.52±0.0195 | 4.69e-21 | 0.523±0.0228 | 8.36e-20 | 0.516±0.0154 | 1.49e-20 | 1.014 |
| Chronos | 0.716±0.0599 | 5.59e-11 | 0.706±0.0587 | 2.35e-8 | 0.686±0.0566 | 1.09e-9 | 1.029 |
| AC-Chronos | **0.501±0.00223** | 8.56e-22 | **0.501±0.00234** | 1.29e-20 | **0.501±0.00194** | 1.2e-21 | **0.9998** |
| Crispy | 0.607±0.0402 | 0.025 | 0.619±0.0426 | 9.72e-4 | 0.588±0.036 | 4.34e-1 | 1.053 |
| GAM | 0.596±0.0389 | 6.85e-4 | 0.608±0.0412 | 1.48e-5 | 0.578±0.0349 | 4.61e-2 | 1.051 |
| Geometric | 0.508±0.0154 | **1.52e-22** | 0.508±0.0177 | **6.54e-21** | 0.507±0.0126 | **9.98e-23** | **1.0002** |
| LDO | 0.621±0.043 | 0.487 | 0.621±0.0436 | 4.65e-1 | 0.611±0.0421 | 5.52e-1 | 1.017 |
| MAGeCK | 0.635±0.0429 | 0.476 | 0.633±0.0419 | 4.62e-1 | 0.625±0.0427 | 5.13e-1 | 1.014 |

**Table S2 - Summary of proximity number bias (mean ± standard deviation) correction across methods in Project Achilles. Results are shown both for genome-wide (GW) and TP53-based proximity bias (i.e. wild-type (WT) and loss-of-function (LOF)). For genome-wide and TP53-based proximity bias, we also performed a *t*-test to assess whether each method's distribution is significantly different from the uncorrected one. For the TP53-based proximity bias, we computed the average ratio (AR) between the WT and LOF proximity bias scores for each method. Bold is the best outcome across methods.**

| Method | GW | P-val | TP53 WT | P-val | TP53 LOF | P-val | AR |
|---|---|---|---|---|---|---|---|

| | | | | | | | |
|---|---|---|---|---|---|---|---|
| Uncorrected | 0.646±0.0821 | NA | 0.643±0.0825 | NA | 0.624±0.0824 | NA | 1.032 |
| CCR | 0.521±0.0371 | 7.89e-13 | 0.522±0.0387 | 8.36e-20 | 0.517±0.0282 | 1.49e-20 | 1.009 |
| Chronos | 0.726±0.0822 | 2.1e-5 | 0.717±0.0809 | 2.35e-8 | 0.69±0.0882 | 1.09e-9 | 1.043 |
| AC-Chronos | **0.501±0.0046** | **1.11e-14** | 0.502±0.0023 | 2.38e-12 | **0.502±0.0056** | 2.66e-14 | 0.9992 |
| Crispy | 0.64±0.0787 | 0.721 | 0.646±0.0768 | 9.72e-4 | 0.609±0.0797 | 4.34e-1 | 1.062 |
| GAM | 0.624±0.0815 | 0.219 | 0.629±0.0814 | 1.48e-5 | 0.595±0.0647 | 4.61e-2 | 1.056 |
| Geometric | 0.52±0.0624 | 8.69e-12 | **0.519±0.062** | **6.54e-21** | 0.52±0.0597 | **9.98e-23** | 0.9979 |
| LDO | 0.643±0.0812 | 0.847 | 0.641±0.0824 | 4.65e-1 | 0.62±0.0837 | 5.52e-1 | 1.035 |
| MAGeCK | 0.595±0.0481 | 6.85e-4 | 0.591±0.045 | 4.62e-1 | 0.591±0.0513 | 5.13e-1 | **1.001** |

**Table S3 - Summary of proximity number bias (mean ± standard deviation) correction across methods in Project Score. Results are shown both for genome-wide (GW) and TP53-based proximity bias (i.e. wild-type (WT) and loss-of-function (LOF)). For genome-wide and TP53-based proximity bias, we also performed a *t*-test to assess whether each method's distribution is significantly different from the uncorrected one. For the TP53-based proximity bias, we computed the average ratio (AR) between the WT and LOF proximity bias scores for each method. Bold is the best outcome across methods.**

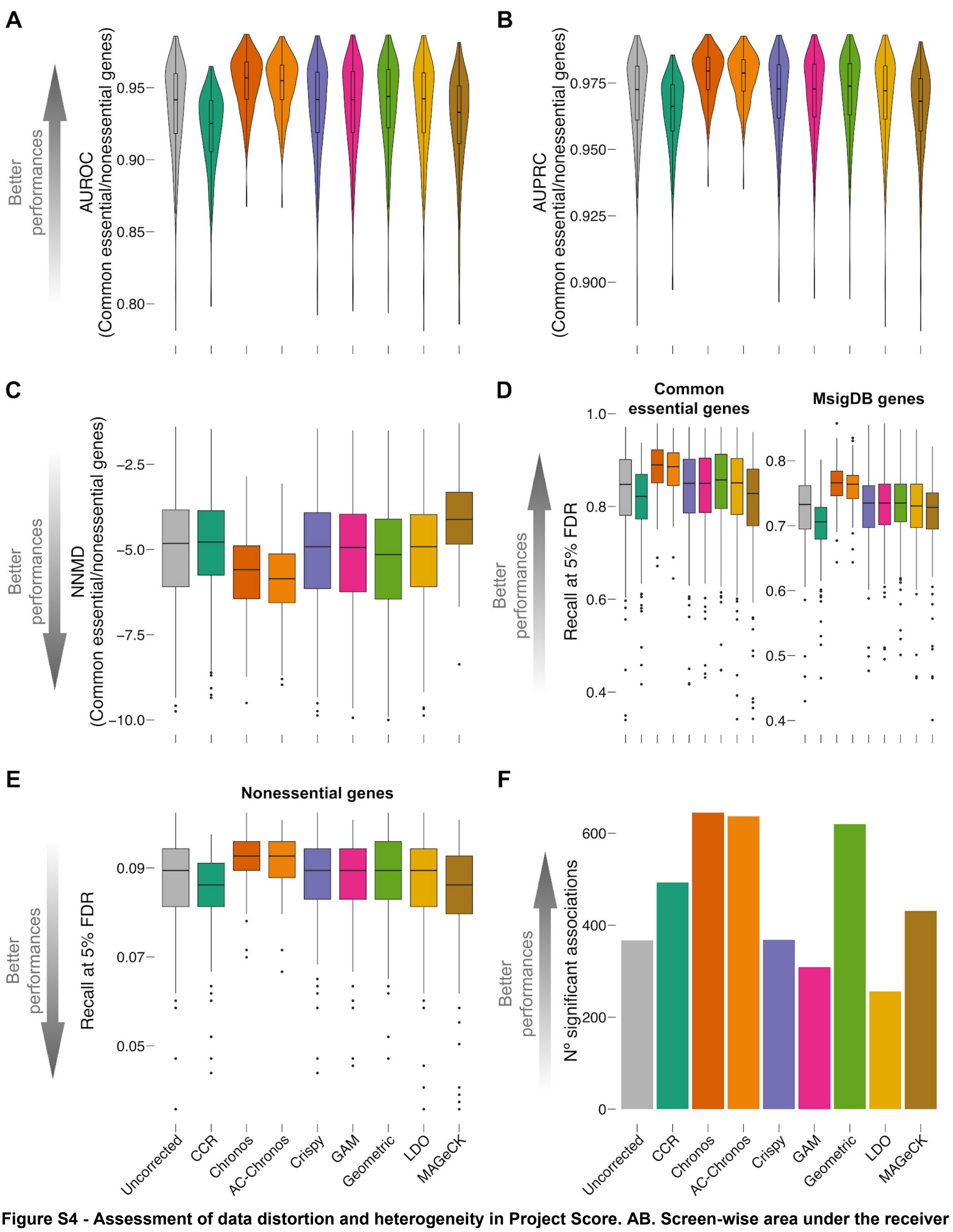


**Figure S4 - Assessment of data distortion and heterogeneity in Project Score. AB. Screen-wise area under the receiver operating characteristic (AUROC) curve (A) and area under the precision-recall curve (AUPRC) (B) were obtained when considering all the screens in Project Score (uncorrected version and processed with all tested methods) as rank-based classifiers of common-essential/non-essential genes, based on the depletion log fold change (LFC). C. Screen-wise null-normalised median difference (NNMD) between the distribution of LFCs of common essential and non-essential log fold change (LFC) distributions for the screens in Project Score (uncorrected version and processed with all tested methods). DE. Screen-wise recall at a 5% false discovery rate was obtained when considering each screen as in ABC but using different gene sets as positive controls (i.e. common essential genes, set of core-fitness genes from MsigDB (E) and non-essential genes (N)) for the Project Score dataset (uncorrected version and processed with all tested methods). F. Number of tissue-specific statistically significant (FDR < 5%) Cancer Functional Events (CFEs) / gene-essentiality associations identified in Project Score (unprocessed version and across correction methods) via a**

systematic two-sided *t*-test. This test assesses differential gene essentiality (as measured by the gene depletion log fold-change (LFC)) across two sub-populations of cell lines stratified based on the presence/absence of the considered CFE.

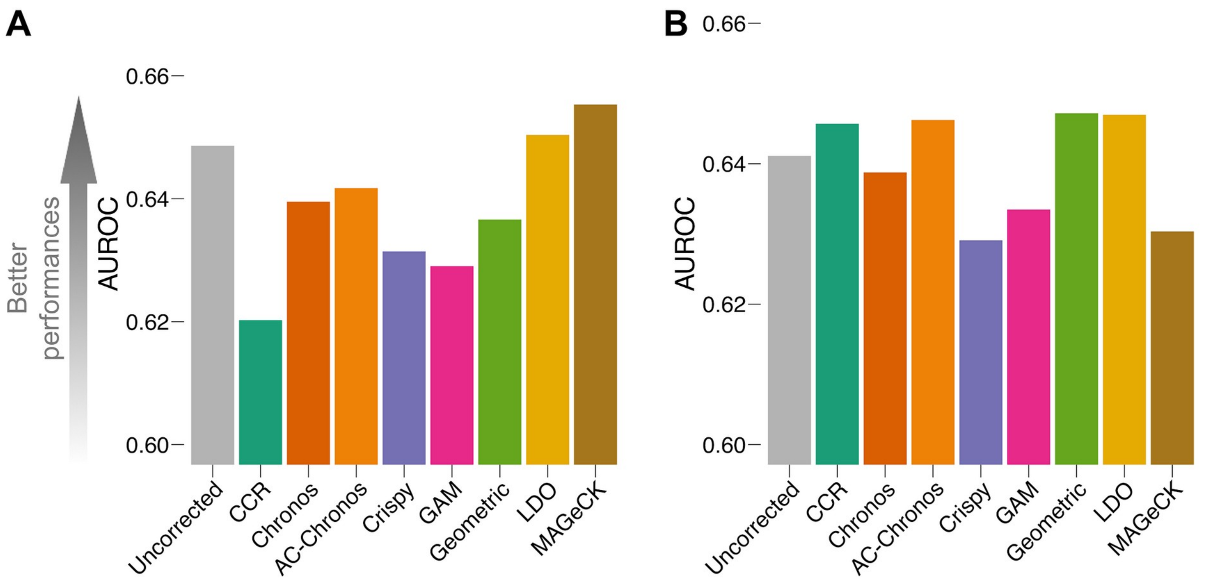


**Figure S5 - Identification of oncogenetic addictions. AB. AUROC obtained when considering all genes' LFCs scaled and pooled across screens in Project Achilles (A) and Project Score (B) (uncorrected version and processed with all tested methods) as a unique rank-based classifier of oncogenic addictions, and using cell line specific mutated oncogenes as positive controls and wild-type unexpressed (TPM < 1) oncogenes as negative controls.**

| Method | AUROC (common essential vs non-essential genes) | AUPRC (common essential vs non-essential genes) | NNMD (common essential vs non-essential genes) | AUROC (oncogenetic addictions) |
|---|---|---|---|---|
| Uncorrected | 0.949±0.0399 | 0.972±0.0239 | -5.19±2.02 | 0.649 |
| CCR | 0.93±0.0357 | 0.963±0.022 | -4.7±1.67 | 0.62 |
| Chronos | **0.961±0.0223** | **0.98±0.0116** | -5.93±1.53 | 0.64 |
| AC-Chronos | 0.96±0.0224 | 0.979±0.0117 | **-6.09±1.59** | 0.642 |
| Crispy | 0.949±0.0398 | 0.972±0.0237 | -5.27±2.06 | 0.631 |
| GAM | 0.949±0.0397 | 0.972±0.0236 | -5.3±2.07 | 0.629 |
| Geometric | 0.949±0.0406 | 0.972±0.0241 | -5.41±2.15 | 0.637 |
| LDO | 0.949±0.0399 | 0.972±0.0239 | -5.22±2.03 | 0.65 |
| MAGeCK | 0.953±0.0356 | 0.974±0.0211 | -5.11±1.8 | **0.655** |

**Table S4 - Summary of recall of common essential and non-essential genes and oncogenetic addictions (mean ± standard deviation) across methods in Project Achilles. Results are based on the following metrics: area under the receiver operating characteristic (AUROC), area under the precision-recall curve (AUPRC), null-normalised median difference (NNMD), and AUROC for oncogenetic addictions. Bold is the best outcome across methods.**

| Method | AUROC (common essential vs non-essential genes) | AUPRC (common essential vs non-essential genes) | NNMD (common essential vs non-essential genes) | AUROC (oncogenetic addictions) |
|---|---|---|---|---|
| Uncorrected | 0.936±0.032 | 0.969±0.0164 | -5.04±1.66 | 0.641 |
| CCR | 0.92±0.0273 | 0.964±0.0141 | -4.91±1.55 | 0.646 |
| Chronos | **0.954±0.0194** | **0.978±0.00902** | -5.74±1.08 | 0.639 |
| AC-Chronos | 0.952±0.0194 | 0.977±0.00904 | **-5.91±1.05** | 0.646 |
| Crispy | 0.937±0.0317 | 0.97±0.0159 | -5.11±1.67 | 0.629 |
| GAM | 0.937±0.0315 | 0.97±0.0158 | -5.15±1.68 | 0.633 |
| Geometric | 0.939±0.0312 | 0.971±0.0156 | -5.32±1.77 | **0.647** |
| LDO | 0.936±0.032 | 0.969±0.0163 | -5.1±1.67 | **0.647** |
| MAGeCK | 0.925±0.0352 | 0.964±0.0181 | -4.12±1.14 | 0.63 |

**Table S5 - Summary of recall of common essential and non-essential genes and oncogenetic addictions (mean ± standard deviation) across methods in Project Score. Results are based on the following metrics: area under the receiver operating characteristic (AUROC), area under the precision-recall curve (AUPRC), null-normalised median difference (NNMD), and AUROC for oncogenetic addictions. Bold is the best outcome across methods.**

| Method | Recall 5% FDR common essential genes | Recall 5% FDR MsigDB genes | Recall 5% FDR non-essential genes |
|---|---|---|---|
| Uncorrected | 0.851±0.133 | 0.692±0.087 | 0.0756±0.0119 |
| CCR | 0.814±0.123 | 0.662±0.0802 | **0.0723±0.011** |
| Chronos | **0.893±0.063** | **0.723±0.0402** | 0.0793±0.00563 |
| AC-Chronos | 0.89±0.063 | 0.72±0.0395 | 0.079±0.00565 |
| Crispy | 0.853±0.131 | 0.695±0.0861 | 0.0758±0.0117 |
| GAM | 0.855±0.13 | 0.696±0.086 | 0.0759±0.0116 |
| Geometric | 0.856±0.132 | 0.694±0.0865 | 0.076±0.0118 |
| LDO | 0.852±0.133 | 0.693±0.0865 | 0.0757±0.0118 |
| MAGeCK | 0.866±0.118 | 0.702±0.0727 | 0.077±0.0105 |

**Table S6 - Summary of recall at 5% false discovery rate of predefined gene sets (i.e. common essential, MsigDB and non-essential genes) and recall curves of amplified and amplified unexpressed genes. Results (mean ± standard deviation) are computed across methods in Project Achilles. Bold is the best outcome across methods. Increased performances are defined as higher recall at 5% FDR for common essential and MsigDB genes, and lower recall at 5% for non-essential genes.**

| Method | Recall 5% FDR common essential genes | Recall 5% FDR MsigDB genes | Recall 5% FDR non-essential genes |
|---|---|---|---|
| Uncorrected | 0.828±0.099 | 0.723±0.0556 | 0.0867±0.0104 |
| CCR | 0.807±0.084 | 0.698±0.0447 | 0.0844±0.00887 |
| Chronos | **0.883±0.0518** | **0.766±0.0297** | 0.0924±0.00549 |
| AC-Chronos | 0.878±0.0517 | 0.762±0.0288 | 0.0919±0.00546 |
| Crispy | 0.832±0.0952 | 0.727±0.0543 | 0.0871±0.01 |
| GAM | 0.833±0.0943 | 0.727±0.0542 | 0.0872±0.00991 |
| Geometric | 0.84±0.0909 | 0.728±0.0518 | 0.0879±0.00958 |
| LDO | 0.83±0.0993 | 0.724±0.0551 | 0.0869±0.0105 |
| MAGeCK | 0.805±0.105 | 0.717±0.0558 | **0.0842±0.0111** |

**Table S7 - Summary of recall at 5% false discovery rate of predefined gene sets (i.e. common essential, MsigDB and non-essential genes) and recall curves of amplified and amplified unexpressed genes. Results (mean ± standard deviation) are computed across methods in Project Score. Bold is the best outcome across methods. Increased performances are defined as higher recall at 5% FDR for common essential and MsigDB genes, and lower recall at 5% for non-essential genes.**

| Method | Project Achilles | Project Score |
|---|---|---|
| Uncorrected | 1740 | 367 |
| CCR | **2240** | 493 |
| Chronos | 1978 | **645** |
| AC-Chronos | 2135 | 637 |
| Crispy | 2116 | 368 |
| GAM | 2179 | 309 |
| Geometric | 1497 | 620 |
| LDO | 2009 | 256 |
| MAGeCK | 1805 | 431 |

**Table S8 - Summary of the total number of significant associations between cancer functional events (CFEs) and strongly selective dependencies across methods in Project Achilles and Project Score. Bold is the best outcome across methods.**
